# Metagenomic and Metatranscriptomic Analysis Reveals Enrichment for Xenobiotic-Degrading Bacterial Specialists and Xenobiotic-Degrading Genes in a Canadian Prairie Two-Cell Biobed System

**DOI:** 10.1101/2021.05.03.441842

**Authors:** J. N. Russell, B. J. Perry, J. Bergsveinson, C. N. Freeman, C. Sheedy, D. Nilsson, L. Braul, C. K. Yost

## Abstract

Biobeds are agriculture-based bioremediation tools used to safely contain and microbially degrade on-farm pesticide waste and rinsate, thereby reducing the negative environmental impacts associated with pesticide use. While these engineered ecosystems demonstrate efficient pesticide removal, their microbiome dynamics remain largely understudied both taxonomically and functionally. As such, further characterization of these parameters may aid in the optimization of biobed management and deployment. This study used metagenomic and metatranscriptomic techniques to characterize the microbial community in a two-cell Canadian biobed system before and after a field season of pesticide application. These culture-independent approaches identified an enrichment of xenobiotic-degrading bacteria, such as *Afipia*, *Sphingopyxis* and *Pseudomonas*, and enrichment and transcription of xenobiotic-degrading genes, such as peroxidases, oxygenases, and hydroxylases, among others; we were able to directly link the transcription of these genes to *Pseudomonas*, *Oligotropha*, *Mesorhizobium*, *Rhodopseudomonas*, and *Stenotrophomonas* taxa.

## Introduction

The agricultural industry benefits from the use of pesticides to protect crops from yield loss due to insects, fungi, and weeds (Aktar *et al*. 2009; Tortella *et al*. 2012). However, point source pollution occurring during the maintenance of pesticide spraying equipment, and the improper disposal of pesticide rinsate and waste (Holmsgaard *et al*. 2017) has polluted soil and aquatic environments at biologically-relevant, toxic concentrations (Chin-Pampillo *et al*. 2015; Spliid *et al*. 2006). To mitigate this problem, on-farm bioremediation ‘biobed’ systems have been developed to collect, biofilter, and degrade on-farm pesticide rinsate and waste using natural soil microflora (Karanasios *et al*. 2010).

Biobeds share a common design, which includes a biologically active biomixture (biomix), which is composed of a 2:1:1 ratio of a lignocellulosic material, peat or compost, and topsoil, and is designed to support a complex pesticide-degrading microbiome (Karanasios *et al*. 2010). Ligninolytic fungi are widely regarded as the primary and initial drivers of biologically-mediated pesticide degradation using lignin-degrading peroxidases and laccases (Bending *et al*. 2002), while bacteria are thought to further metabolize those degradation products (Pinto *et al*. 2016) using mono- and dioxygenases, hydroxylases (Uotila *et al*. 1992), or lyases, etc. (Singh *et al*. 1999). However, there has been a limited number of studies conducted to describe the nature and structure of these microbial communities, with studies arriving at divergent conclusions regarding the effect pesticides have on diversity and community membership (Bergsveinson *et al*. 2018; Holmsgaard *et al*. 2017; Milanovic *et al*. 2010; Coppola *et al*. 2011; Góngora-Echeverría *et al*. 2018; Tortella *et al*. 2013).

This study aimed to elucidate the resident microbial communities within a two-cell Canadian biobed system, and their associated functionality through metagenomic and metatranscriptomic analyses. By sampling before and after a season of pesticide use, we measured how the communities shifted both taxonomically and functionally in response to pesticide exposure, and used comparative approaches to measure abundances and diversity of aromatic-degrading genes.

## Results

### Metagenomic and Metatranscriptomic Sequencing

12-25 million DNA sequencing reads remained per metagenomic sample after quality filtering, with an average of 10.19 % of reads being removed. The metatranscriptomic dataset included 60-90 million reads per sample after quality filtering, with an average of 4.85 % of reads being removed. Nonpareil analysis estimated that the metagenomic dataset achieved ~10-40 % coverage of the total community (Figure S3), while the metatranscriptomic dataset achieved ~75-85 % (Figure S4). MetaPhlAn2 aligned ~99.96 % of metagenomic reads to bacterial genomes and ~0.04 % to viral genomes; the metatranscriptomic dataset aligned ~ 83.32 % of reads to bacterial genomes, ~16.47 % to viral genomes, and ~0.21 % to eukaryotic genomes. The HUMAnN2 pipeline aligned ~ 7.1 % of reads to gene families for the metagenome, and ~11.01 % for the metatranscriptome; alignment was increased by ~ 40 % when using the less stringent Uniref50 database, although we chose to use the Uniref90 results for data analysis. We additionally mapped the metagenome to Kraken 2’s fungal-specific RefSeq database (v2.1.1; Wood *et al*. 2019); this database was further customized through the addition of notable lignin-degrading fungal genomes (*Aspergillus niger*, *Aspergillus nidulans*, *Ganoderma lucidum*, *Irpex lacteus*, *Phanerochaete crysosporium*, *Pleurotus ostreatus*, *Trametes* sp., and *Trichoderma oligosporum*). ~ 0.05 % of these reads aligned to this fungal database, suggesting that these metagenomic samples were dominated by bacterial DNA.

### Diversity and Community Similarity

Biobed 1 pre-pesticide (BB1.Pre) had 55 genera and 63 species, Biobed 1 post-pesticide (BB1.Post) had 58 genera and 81 species, Biobed 2 pre-pesticide (BB2.Pre) had 43 genera and 46 species, and Biobed 2 post-pesticide (BB2.Post) had 33 genera and 36 species (phyla, class, and order diversity is described in Figure S5). While there was a decrease in beta diversity in Biobed 1 after pesticide application, no statistical difference in the community diversity following pesticide application for either biobed cell was observed (Figure 1A).

**Figure 1:**
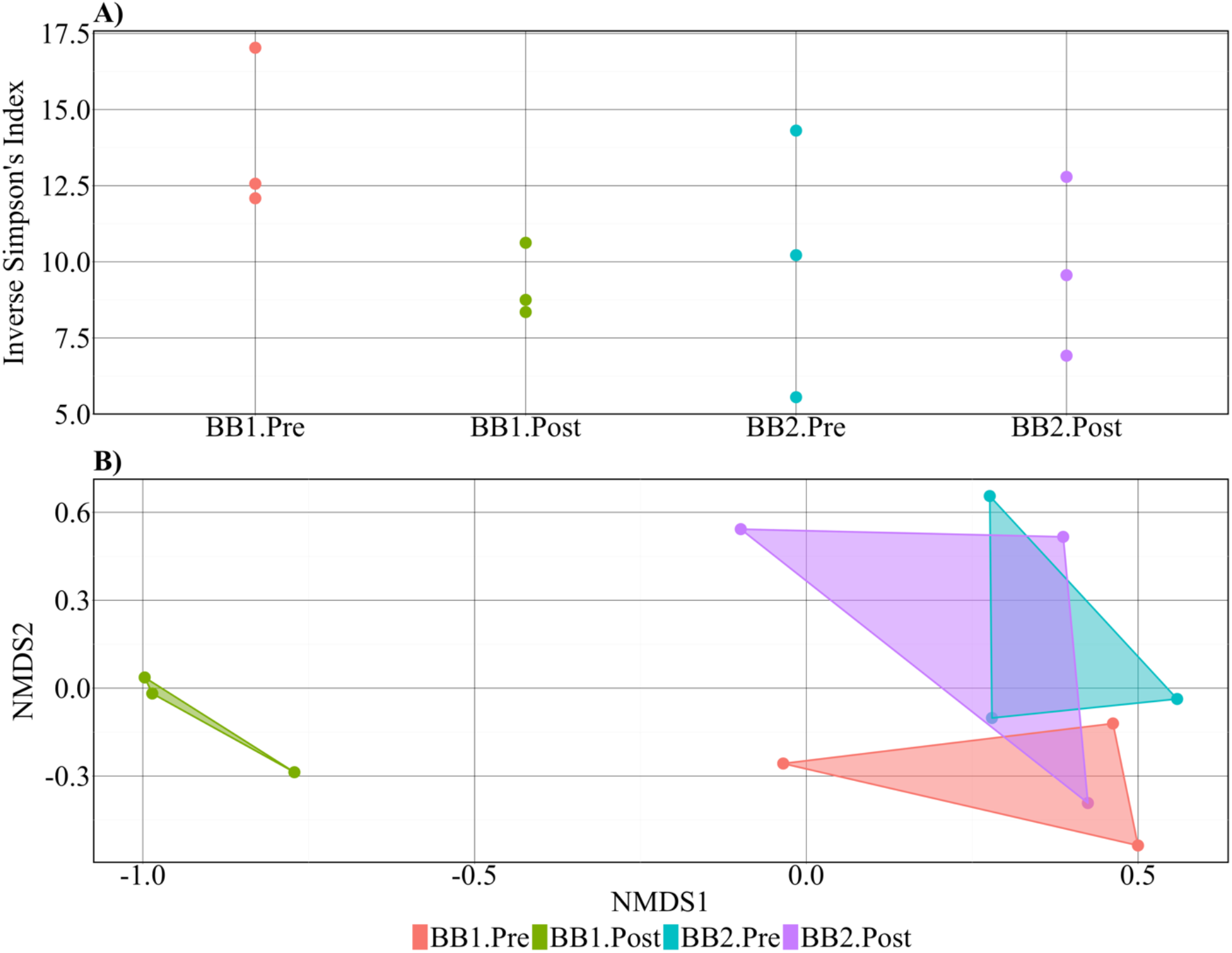
**A)** Beta diversity analysis and **B)** Non-Metric Multidimensional Scaling (NMDS) analysis for biobed microbial communities at the level of species. *BB(Biobed), *Pre - pre-pesticide application, *Post – post-pesticide application.

The spatial separation of biobed samples within the NMDS space (Figure 1B) demonstrates that there is a shift in community membership in Biobed 1 after pesticide application, but not Biobed 2. Additionally, BB1.Pre has a community structure that is more similar to Biobed 2 than to BB1.Post, indicating that both biobeds started with relatively similar communities, and the community in Biobed 1 changed after the season of pesticide application.

LEfSe analysis identified 113 discriminatory taxa (LDA Score > |2.0|) between BB1.Pre and BB1.Post, 24 of which were at the genus level (Figure 2B). There were only 2 differences between BB2.Pre and BB2.Post: BB2.Pre showed significantly higher abundance of the family *Comamonadaceae* (LDA = 4.13), and BB2.Post showed significantly higher abundance of the order Burkholderiales (LDA = 4.22). From these taxa, *Mesorhizobium*, *Brevundimonas*, *Leifsonia*, *Phenylobacterium*, *Pseudoxanthomonas*, *Variovorax*, *Agromyces*, and *Caulobacter* are present at ≥ 0.01 relative abundance in BB1.Pre, and *Sphingopyxis*, *Pseudomonas*, *Afipia*, *Hyphomicrobium*, *Sphingobium*, *Ochrobactrum*, and *Oligotropha* are present at ≥ 0.01 relative abundance in BB1.Post (Figure 2A).

**Figure 2:**
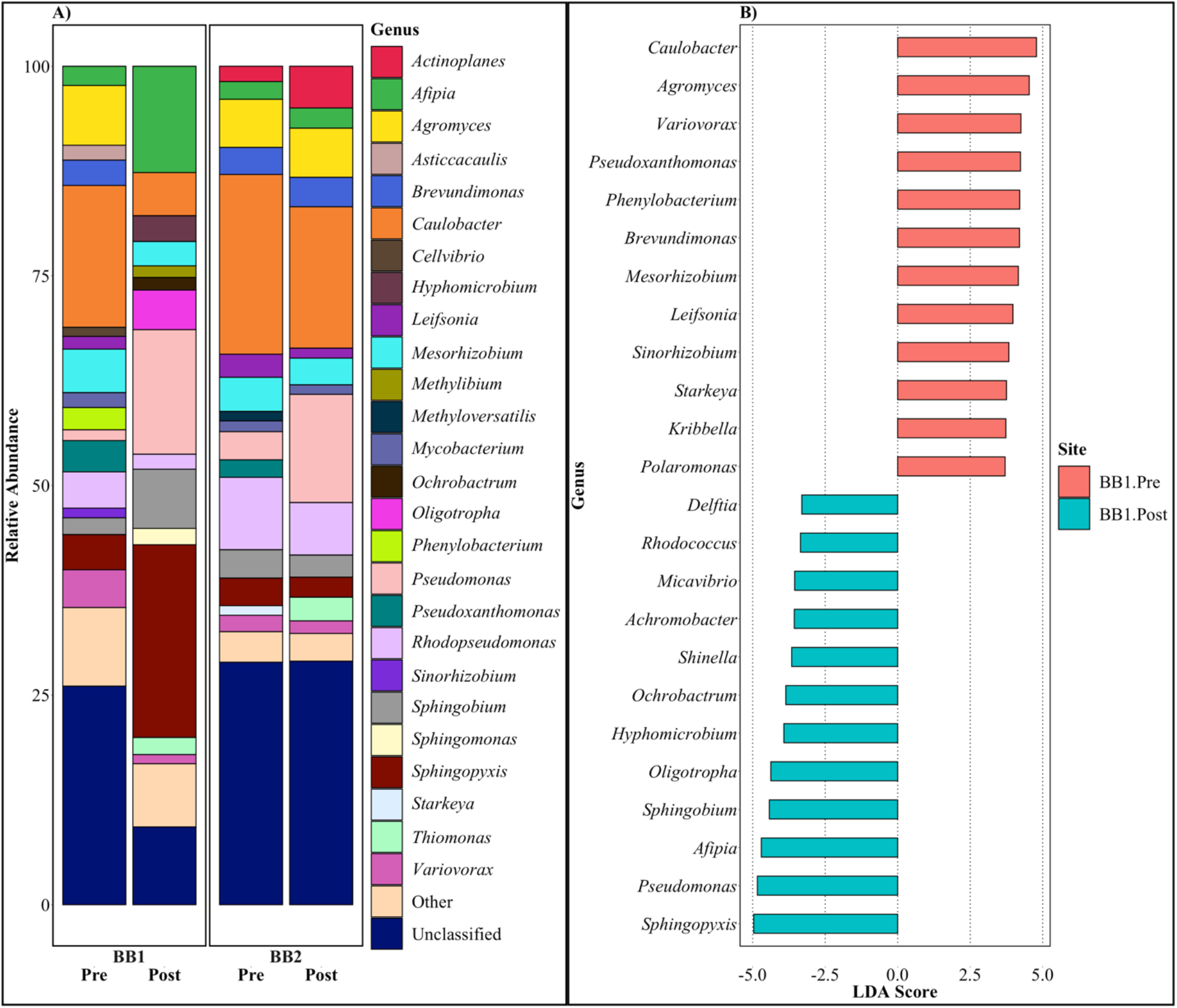
**A)** The relative abundance of genera which contributed an average relative abundance ≥ 0.01; ‘Other’ refers to taxa were present at < 0.01 relative abundance, and ‘Unclassified’ refers to taxa that were not classifiable at the level of genera. Values were averaged between the three sampling replicates. B) The differentially abundant genera (LDA Score > |2.0|) between BB1.Pre and BB1.Post, as determined through LDA effect size using LEfSe. *BB(Biobed), *Pre - pre-pesticide application, *Post - post-pesticide application.

### Functional Genomics

Kruskall-Wallis analyses demonstrated no significant differences in gene abundance following pesticide application in Biobed 2, which reflects the data from the taxonomic analyses (Figure 1B). However, within Biobed 1, there are 1.11-11.8-fold increases in the gene abundance of peroxidases, monooxygenases, lyases, hydroxylases, dioxygenases, dehalogenases, cytochrome P450, and amidases, and 2.87-18.27-fold increases in gene abundance, and transcription of genes related to the degradation of phenols, hydrocarbons, catechol, biphenyls, antibiotics, and non-specific aromatic structures in Biobed 1, as well as transcription of these genes (Figure 3). Laccases show limited gene abundance across the biobed samples, and transcription only in BB1.Post. Similarly, hydrogenases show an increase in gene abundance but not transcription in BB1.Post.

**Figure 3:**
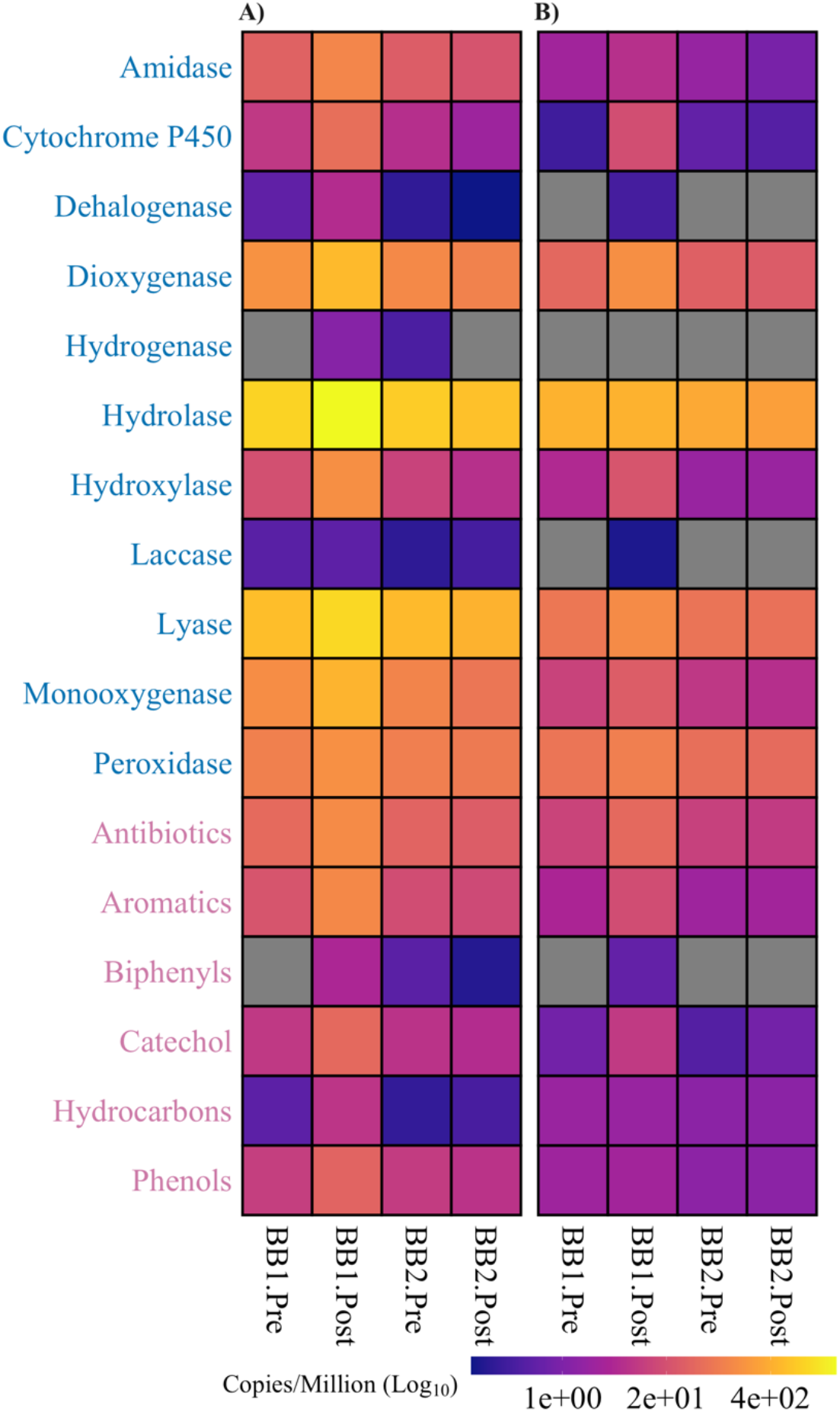
The abundance in copies per million of genes related to aromatic structure degradation observed in the A) metagenome and B) metatranscriptome; keywords in blue represent enzymatic categories, and keywords in pink represent degradation targets. Genes that showed significantly different abundance across biobed conditions through a Kruskal-Wallis statistical test were identified and summarized into the listed broad categories. Grey indicates 0 gene abundance for that site. *BB (Biobed), *Pre - pre-pesticide application, *Post - post-pesticide application.

From the taxa highlighted in Figure 2, the genes of interest (Figure 3) were metagenomically linked to *Variovorax*, *Starkeya*, *Sphingomonas*, *Sphingobium*, *Sphingopyxis*, *Shinella*, *Rhodopseudomonas*, *Rhodococcus*, *Pseudomonas*, *Phenylobacterium*, *Oligotropha*, *Ochrobactrum*, *Mycobacterium*, *Micavibrio*, *Methyloversatilis*, *Methylibium*, *Mesorhizobium*, *Hyphomicrobium*, *Delftia*, *Afipia*, and *Achromobacter*, *and metatranscriptomically linked to Caulobacter*, *Hyphomicrobium*, *Pseudomonas*, *Mesorhizobium*, *Micavibrio*, *Oligotropha*, *Sphingobium*, and *Rhodopseudomonas* in Biobed 1; the majority of reads were associated with unclassified genomes (Figure S6). It should be noted that *Stenotrophomonas*, which was not highlighted in Figure 2, was also shown to substantially contribute to the transcription of several aromatic-degrading genes (> 5.0 %), most notably laccases and antibiotic-degrading genes.

## Discussion

The purpose of this study was to measure how pesticide application impacts microbial community membership and gene abundances in a two-cell biobed system in Lethbridge, Alberta, Canada. The present findings agree with other studies that have found pesticide exposure within a biobed environment significantly shifts the community structure and membership but does not significantly change the diversity of the microbial community (Figure 1). Specifically, after a season of pesticide treatment, there is a significant decrease in soil-associated microbes, such as *Caulobacter* (Wilhelm. 2018), *Agromyces* (Gledhill & Casida. 1969), *Brevundimonas* (Yoon *et al*. 2007), and *Mesorhizobium* (Wang *et al*. 2002), and an increase in microbes known to degrade various types of xenobiotics, such as *Sphingopyxis* (Wang *et al*. 2010; Aranda *et al*. 2003; Yamatsu *et al*. 2006; Kertesz & Kawasaki. 2010; Chen *et al*. 2013), *Afipia* (Isaka *et al*. 2016; Zhang *et al*. 2009), and *Pseudomonas* (Nishino & Spain. 1993; Zylstra *et al*. 1988) (Figure 2B). These results demonstrate a shift towards a xenobiotic-degrading specialist community, which is evidence towards the key players in biobed-based pesticide degradation. However, this biobed system differs from the microbiomes of four other Canadian Prairie biobed systems (Bergsveinson *et al*. 2018), which had a shared core microbiome mostly consisting of Actinomycetales, Acidobacteria, Bacteroidetes, Rhizobiales, and Sphingobacteriales; the most abundant orders in the Lethbridge biobed post-pesticide treatment were Sphingomonadales, Rhizobiales, and Pseudomonadales (Figure S5). This suggests that biobed microbiomes can be unique, even regionally, and that a true core microbiome may not exist, which could be due to biomix sources, pesticide variety, environmental conditions, or functional overlap between different microbes.

Our metagenomic and metatranscriptomic analyses support the idea that pesticides are degraded in biobeds through a complex network of biological activity, as Biobed 1 demonstrated an increase in abundance of several genes related to aromatic degradation, such as peroxidases, monooxygenases, and cytochrome P450, etc., and the degradation of specific xenobiotics (Figure 3). However, the transcriptional activity of these genes can only be taxonomically linked to a small subset of the most abundant/significantly shifting bacteria: *Pseudomonas*, *Sphingobium*, and *Oligotropha* (Figure S6). Interestingly, we also linked post-pesticide transcription to two genera that decrease in abundance post-pesticide treatment: *Mesorhizobium* and *Rhodopseudomonas. Stenotrophomonas*, a genus that is also known to degrade xenobiotics (Ryan *et al*. 2009), also showed notable transcription despite being present at < 0.01 relative abundance (Figure 2). This indicates that low abundance microbes may remain an active part in the degradation of pesticides, and that high taxonomic abundance may not directly relate to increased functional activity. However, the majority of mapped reads were associated with ‘unclassified’ genomes, which demonstrates that a large component of this biological network remains understudied and/or that some of the genes involved in pesticide degradation are not taxa specific. This observation prompts the need for exploring biobed mechanics and soil microbiomes more thoroughly, perhaps through culture-based approaches, to more completely characterize the genomes of pesticide-degrading bacteria.

Finally, the absence of fungal representation in this study is notable, as the literature suggests that ligninolytic fungi are the primary drivers of pesticide metabolism in biobeds (Pinto *et al*. 2016), and biomixes are formulated to specifically encourage their growth (Castillo *et al*. 2007). It is possible that the conditions of this biobed system did not encourage fungal growth, that fungi were inhibited by fungicides, or that the fungal community did not have sufficient time to establish. Despite this, this study provides evidence that specific bacteria, such as *Pseudomonas* or *Stenotrophomonas* species, might be key players in pesticide degradation within newly constructed biobed systems, and future studies could explore the temporal changes that occur to the biobed as the system ages.

## Experimental Procedures

### Biobed Construction and Operation

The biobed used for this study was built by Agriculture and Agrifood Canada (AAFC) at the Lethbridge Research and Development Center in Lethbridge, Alberta, Canada; details on construction and operation can be found in the supplemental methods section.

### Biomix Sample Collection and Nucleic Acid Preparation

Three replicate biomix samples were collected for metagenomic and metatranscriptomic profiling from each biobed at the beginning of the operational field season before pesticide application (May 2016), and at the end of the field season (October 2016). These samples were taken from a fixed triangular pattern approximately 2.0 m apart, and 10.0 cm in depth after removal of the top 1.0 cm of biomix. Pre-pesticide samples were collected prior to grass seeding, and that post-pesticide samples were taken from the same locations, in between plant rows. Samples were collected in sterile 50 mL conical tubes and were immediately snap-frozen in liquid nitrogen, transported on dry ice, and stored at −80 °C.

Samples were homogenized and nucleic acids were extracted using the MoBio RNA PowerSoil^®^ Total RNA Isolation Kit (Qiagen, CA, USA) and the MoBio RNA PowerSoil^®^ DNA Elution Accessory Kit (Qiagen, CA, USA) from the same 2.0 g of sampled material. RNA was cleaned using a Qiagen RNeasy Min-Elute Cleanup Kit (Qiagen, CA, USA), residual phenol was removed using chloroform (3:1), and samples were DNase-treated using the ThermoFisher TURBO DNA-*free*^™^ Kit (ThermoFisher Scientific, ON, CA). RNA concentrations were established through a Qubit dsRNA HS assay (ThermoFisher Scientific, ON, CA), and RNA integrity was established through an Agilent RNA assay (Agilent, CA, USA). DNA was quantified using a Qubit BR dsDNA assay (ThermoFisher Scientific, ON, CA), and quality was assessed through agarose gel electrophoresis (2 %).

### Metagenomic and Metatranscriptomic Sequencing

Metagenomic sequencing libraries were constructed by Genome Quebec using an Illumina TruSeq DNA PCR-Free Library Preparation kit (Illumina, Inc., CA, US) with a final size selection of 400 bp. All 12 libraries were multiplexed into a single Illumina HiSeq 4000 (Illumina, Inc., CA, US) lane for 100 bp paired-end sequencing. For metatranscriptomic sequencing, the three RNA samples from pre- and post-pesticide applications were pooled together for each biobed, for a total of four samples (Biobed 1 and 2, pre- and post-pesticide application). Metatranscriptomic sequencing libraries were also constructed by Genome Quebec using an Illumina TruSeq RNA Library Preparation kit (Illumina, Inc., CA, US) after rRNA depletion, with both bacterial and fungal Illumina Ribo-Zero rRNA Depletion Kits (Illumina, Inc., CA, US). All four libraries were sequenced on a single Illumina HiSeq 4000 lane for 100 bp paired-end sequencing.

### Data Analysis

Metagenomic and metatranscriptomic datasets were inspected using FastQC (v0.11.8; Andrews. 2014) and MultiQC (v1.7; Ewels *et al*. 2016), and trimmed and quality filtered (Q30) using Trimmomatic (v.0.38; Bolger *et al*. 2014) with a minimum length of 60 bp, trimming of the first 5 bp and the last 30 bp, and trimming of reads whose average quality dropped below 30 over a 10 bp window. Sequencing coverage was determined using Nonpareil (v.3.3.3; Rodriguez-R *et al*. 2018) with the kmer algorithm. Taxonomic identification was conducted through MetaPhlAn2 (v2.7.8; Truong *et al*. 2015) and functional gene identification was conducted through HUMAnN2 (v0.11.2; Franzosa *et al*. 2018) using the ChocoPhlAn and Uniref90 databases. All data visualization was plotted in R (R Core Team. 2020), using the following R packages: Phyloseq (v 1.22.3; McMurdie & Holmes. 2013), ggplot2 (v.3.2.1; Wickham. 2016), viridis (v.0.5.1; Garnier. 2018), ggthemes (v.4.2.0; Arnold. 2019), scales (v.1.1.0; Wickham & Seidel. 2019), reshape2 (v.1.4.3; Wickham. 2007), patchwork (v.1.0.0; Pedersen. 2019), gridExtra (v.2.3; Augie. 2017), tidyr (v.1.0.2; Wickham & Henry. 2020), ggpubr (v.0.2.5; Kassambara. 2020), and Rmisc (v.1.5; Hope. 2013).

### Statistical Analysis

Community diversity of the metagenome was calculated using the Inverse Simpson’s Diversity Index, and further statistically compared via ANOVA and Tukey’s Honestly Significant Difference tests (P > 0.05) in R. Differences in biobed community membership pre- and post-pesticide application were compared using non-metric multidimensional scaling (NMDS) tests. Both Inverse Simpson’s Diversity and NMDS analyses were performed using Phyloseq (v 1.22.3; McMurdie & Holmes. 2013). LEfSe (Galaxy Version 1.0; Segata *et al*. 2011) was used to identify which taxa were responsible for statistically significant differences between communities; LDA effect sizes over 2.0 are considered significant (P > 0.05), as specified by the developer. Lastly, to identify the genes exhibiting significantly different abundances across the biobed system, the Kruskal-Wallis test (P > 0.05) was used on the HUMAnN2 ‘gene families’ output (normalized to copies per million). This list of significant genes was searched for genes that code for aromatic- and xenobiotic-degrading enzymes, and the gene abundance data from these categories were mathematically summed for broad comparison; specific keywords can be found in Figure 3, but also included ‘phenoloxidase’, ‘phosphatase’, ‘polycyclic’, ‘trichloroethylene’, ‘chlorophenol’, and ‘pesticide’, although these keywords had 0 gene abundance. Finally, the metatranscriptomic dataset, which only had one sample per biobed timepoint, was not used for statistical analysis, but was used to corroborate the metagenomic analysis.

## Acknowledgments

We would like to acknowledge Jim Daschle for his construction and maintenance of the biobeds, and the field staff of the Lethbridge Research and Development Center for operation of the biobeds.

