## Supporting Information for "Metagenomic and Metatranscriptomic Analysis Reveals Enrichment for Xenobiotic-Degrading Bacterial Specialists and Xenobiotic-Degrading Genes in a Canadian Prairie Two-Cell Biobed System"

***Supplemental Experimental Procedures***

*Biobed Construction and Operation*

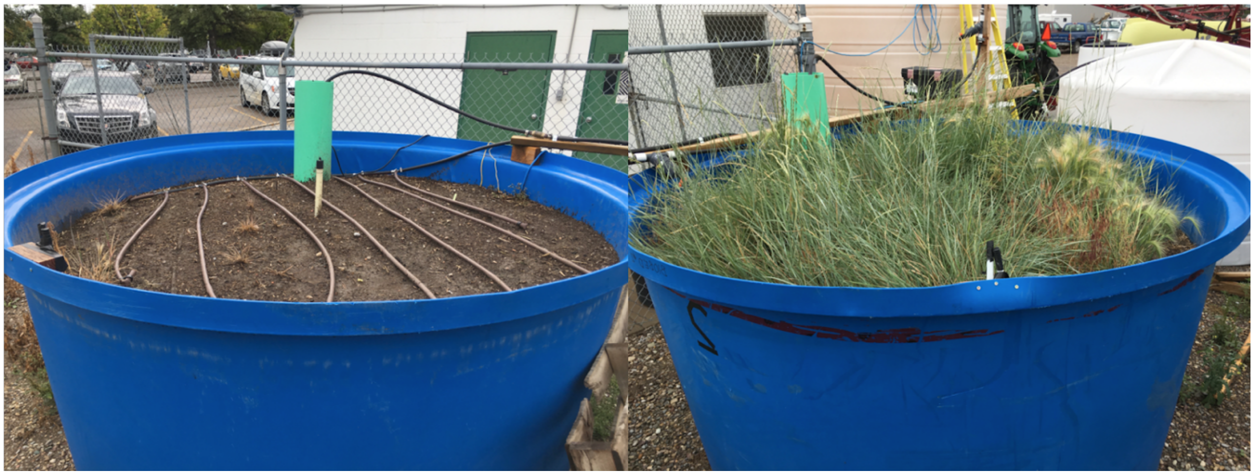

**Figure S1:** Biobed 1 (left) and Biobed 2 (right) at the end of the field season (October 2016).

The biobed was constructed as a two-cell system, whereby the pesticide waste was applied over two sequential biobeds. Pesticide rinsate was first collected on a collection pad, pumped into a holding tank, and applied over the primary biobed (Biobed 1) (20 L/day). The flow-through was pumped from the bottom of Biobed 1 and re-applied over a secondary biobed (Biobed 2) (20 L/day). The final effluent was collected in a holding tank and discharged on a pesticide rinsate-dedicated septic field. The biobeds consisted of above-ground plastic containers that were 2.44 m in diameter and 1.27 m in height with 0.15 m of crushed rock at the bottom. The biomix, which was composed of a 2:1:1 ratio of chopped straw, compost, and topsoil from plots which had previously been exposed to pesticides during agricultural farming, was composted *in situ* for 2 months prior to transfer into the biobeds. This composting process included the periodic addition of water to maintain moisture, as well as physically turning the biomix every two weeks. Both biobeds were seeded with grass after pre-pesticide samples were collected, but only Biobed 2 experienced grass establishment by the end of the field season (Figure S1).

During the sampling period of June to September 2016, 10 fungicides, 24 herbicides, 6 insecticides, and 1 nematicide (Table S1) were applied to the biobed system in an undefined mixture of pesticide rinsate, sourced from pulse crop pesticide applications. Pesticide concentrations were measured monthly from the influent and the effluents of both biobeds using methods described in Sheedy *et al.* (2019); mean pesticide reductions can be found in Figure S2.

*Supplemental Tables and Figures*

| Pesticide Type | Pesticide | Influent (µg/L) | BB1 Effluent (µg/L) | BB2 Effluent (µg/L) |
|---|---|---|---|---|
| **Fungicide** | Azoxystrobin | 0.3295 | 0.0699 | 0.0516 |
| | Boscalid | 6.5856 | 0.0050 | 0.0000 |
| | Difenoconazole I | 40.8026 | 0.0000 | 0.0000 |
| | Ethalfluralin | 4.5771 | 0.0000 | 0.0000 |
| | Fludioxonil | 2.1632 | 0.0000 | 0.0000 |
| | Metalaxyl | 17.2431 | 9.8230 | 6.7792 |
| | Metconazole | 48.7181 | 0.0000 | 0.0000 |
| | Propiconazole I | 10.7937 | 0.0000 | 0.0000 |
| | Prothioconazole-desthio | 15.2448 | 0.0110 | 0.0000 |
| | Pyraclostrobin | 80.7989 | 0.0000 | 0.0000 |
| | Tebuconazole | 74.7015 | 0.0000 | 0.0066 |
| **Herbicide** | 2,4-D * | 3805.3340 | 1.7938 | 0.0714 |
| | 2,4-DB ** | 3.8251 | 0.0000 | 0.0000 |
| | Atrazine | 0.1566 | 0.0000 | 0.0000 |
| | Bentazone | 3577.7941 | 3023.2332 | 1955.7327 |
| | Bromoxynil | 3640.5737 | 0.1251 | 0.1088 |
| | Carfentrazone-ethyl | 0.4610 | 0.0000 | 0.0000 |
| | Clopyralid | 496.9074 | 406.4409 | 428.1238 |
| | Dicamba | 556.7244 | 134.0042 | 34.4112 |
| | Dichlorprop | 2.3632 | 0.0057 | 0.0000 |
| | Diclofop | 7.0215 | 0.0422 | 0.0000 |
| | EPTC *** | 31.6065 | 0.1490 | 0.0000 |

| | | | | |
|---|---|---|---|---|
| | Ethalfluralin | 1.3821 | 0.0000 | 0.0000 |
| | Ethofumesate | 0.1985 | 0.0000 | 0.0000 |
| | Fenoxaprop | 2.3340 | 0.0000 | 0.0000 |
| | Fluroxypyr | 418.2473 | 27.3176 | 0.7162 |
| | Hexazinone | 5.2216 | 0.4885 | 0.1115 |
| | Imazethapyr | 2065.9237 | 1162.7101 | 101.6695 |
| | MCPA **** | 8943.5325 | 7.8875 | 0.2647 |
| | Mecoprop | 1707.9129 | 13.3367 | 0.0747 |
| | Metolachlor | 0.0546 | 0.0000 | 0.0000 |
| | Quizalofop-ethyl | 0.0863 | 0.0000 | 0.0000 |
| | Simazine | 0.2351 | 0.0000 | 0.0000 |
| | Triallate | 435.5562 | 0.0016 | 0.0043 |
| | Trifluralin | 0.4004 | 0.0000 | 0.0000 |
| **Insecticide** | Chlorpyrifos | 0.3631 | 0.0000 | 0.0000 |
| | Cyhalothrin-lambda | 14.1882 | 0.0000 | 0.0000 |
| | Deltamethrin | 0.0957 | 0.0000 | 0.0000 |
| | Malathion | 0.2723 | 0.0000 | 0.0000 |
| | Spiromesifen | 0.2315 | 0.0000 | 0.0000 |
| **Nematicide** | Dichlofenthion | 0.0337 | 0.0000 | 0.0000 |

**Table S1:** The average pesticide concentrations (µg/L) detected in the influent and biobed effluents from measurements taken between June 15th, 2016 and September 30th, 2016. *2,4-D (2,4-Dichlorophenoxyacetic Acid), **2,4-DB (4-(2,4-Dichlorophenoxy)butyric Acid), *** EPTC (S-Ethyl Dipropylthiocarbamate), ****MCPA (2-Methyl-4-Chlorophenoxyacetic Acid).

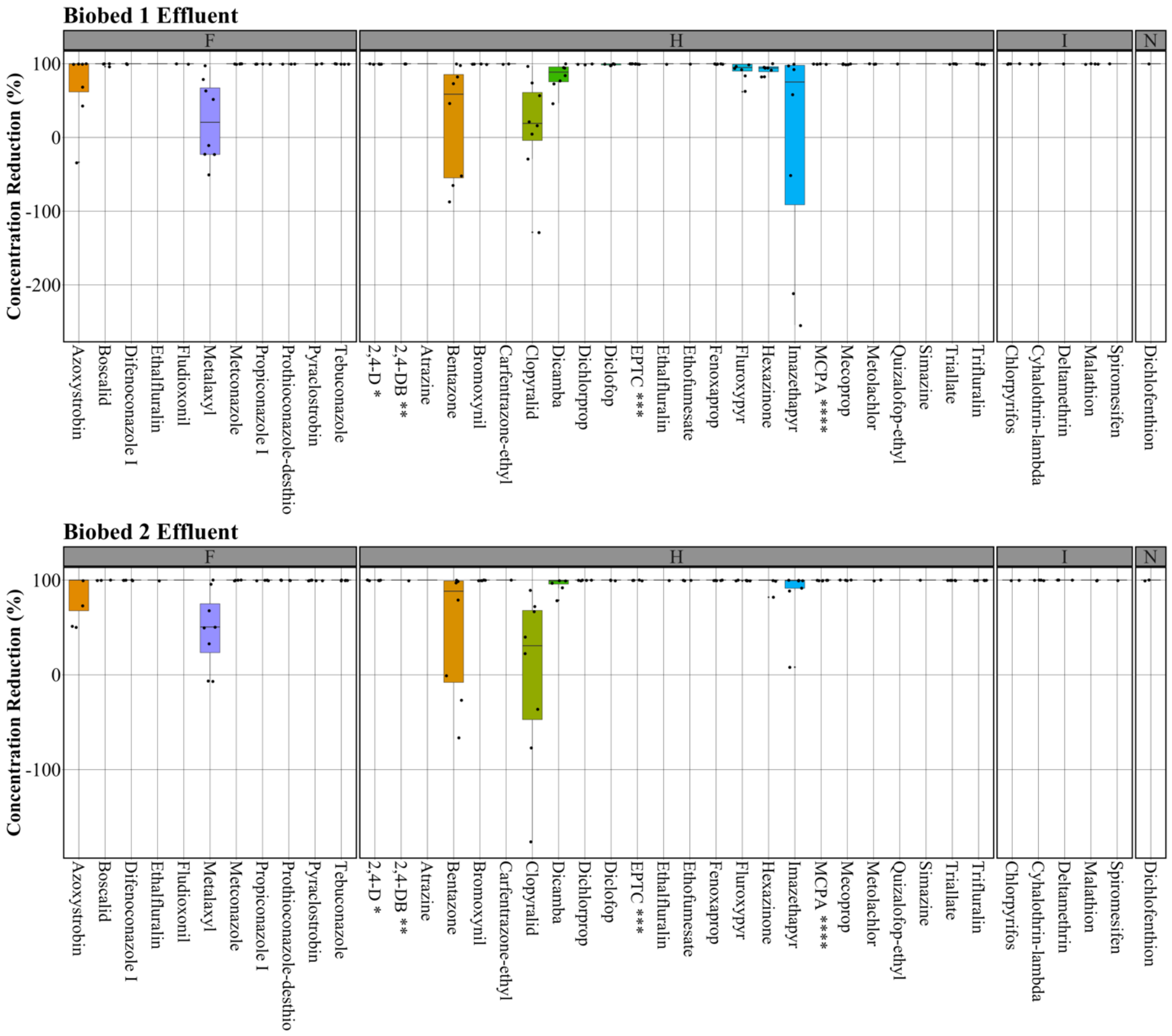


**Figure S2:** The concentration reduction (%) of fungicides (F), herbicides (H), insecticides (I), and nematicides (N) after Biobed 1 and Biobed 2 treatment, compared to initial influent concentrations. True concentration averages, as measured monthly between June and September 2016, can be found in Table S1; this data was measured and provided by the researchers at AAFC. Negative reductions indicate higher concentrations in the effluent than the influent.

*2,4-D (2,4-Dichlorophenoxyacetic Acid), **2,4-DB (4-(2,4-Dichlorophenoxy)butyric Acid), *** EPTC (S-Ethyl Dipropylthiocarbamate), ****MCPA (2-Methyl-4-Chlorophenoxyacetic

Acid). There was an overall significant reduction in pesticide concentration of $\geq 90.20$ % for all four categories of pesticides, as confirmed through ANOVA ($P < 0.05$).

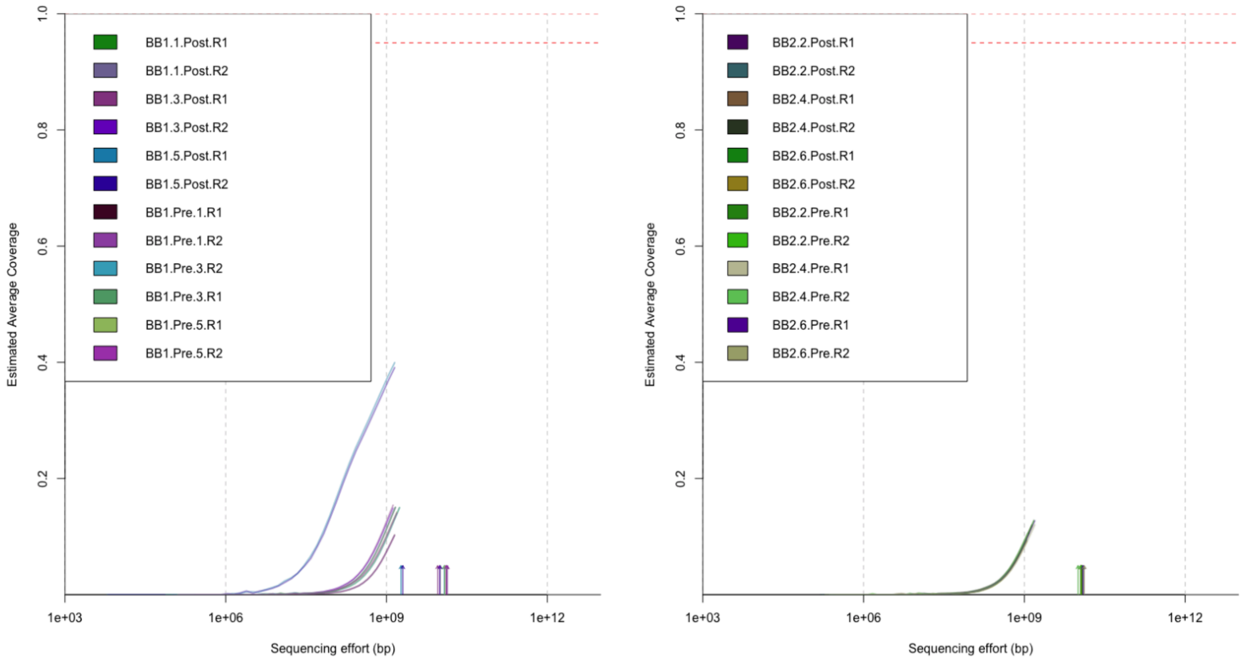


**Figure S3:** The estimated average coverage for the metagenomic dataset for Biobed 1 (left) and Biobed 2 (right), as calculated by Nonpareil (v.3.3.3) (Rodriguez-R *et al.* 2018).

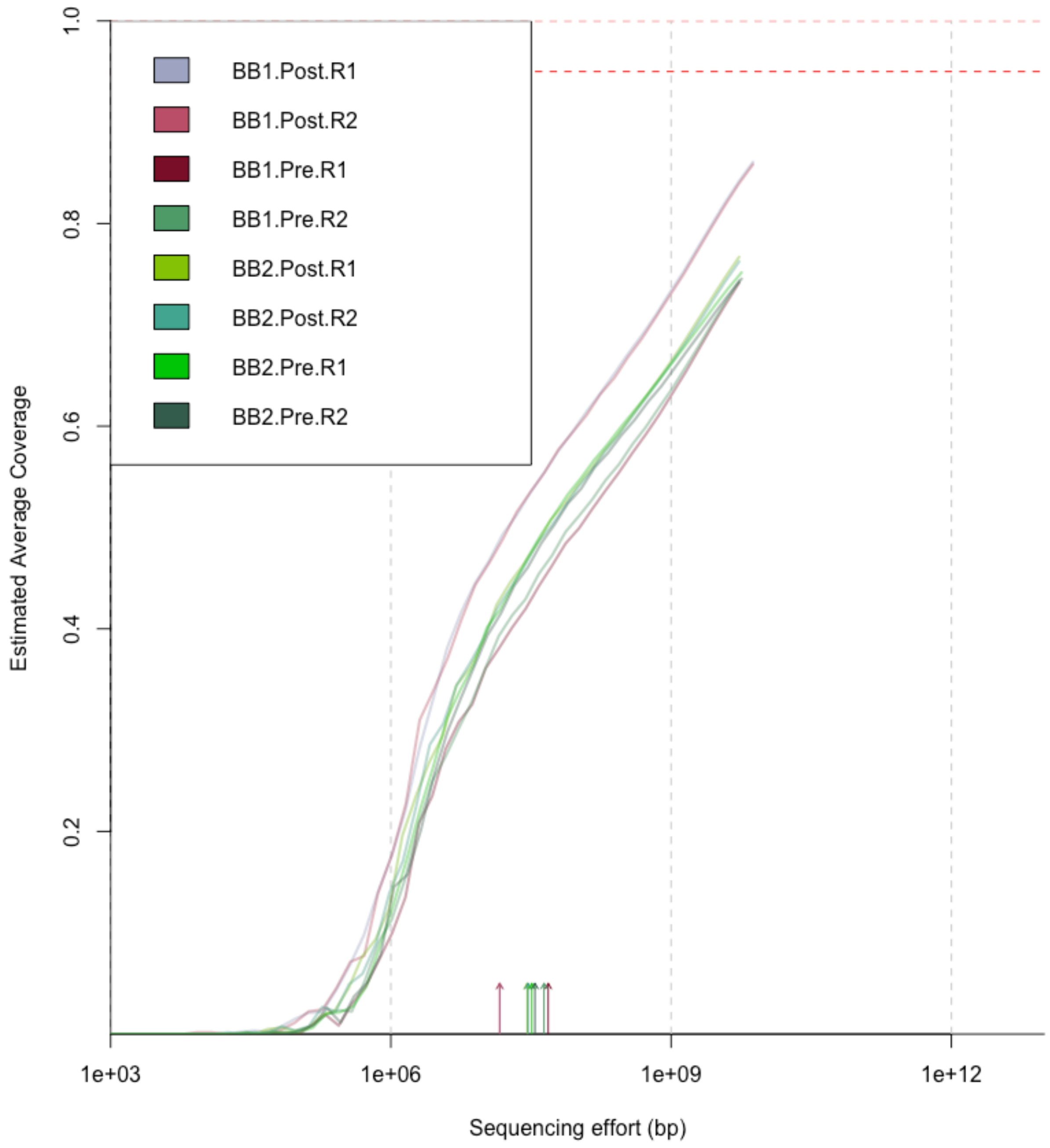


**Figure S4:** The estimated average coverage for the metatranscriptomic dataset, as calculated by Nonpareil (v.3.3.3) (Rodriguez-R *et al.* 2018).

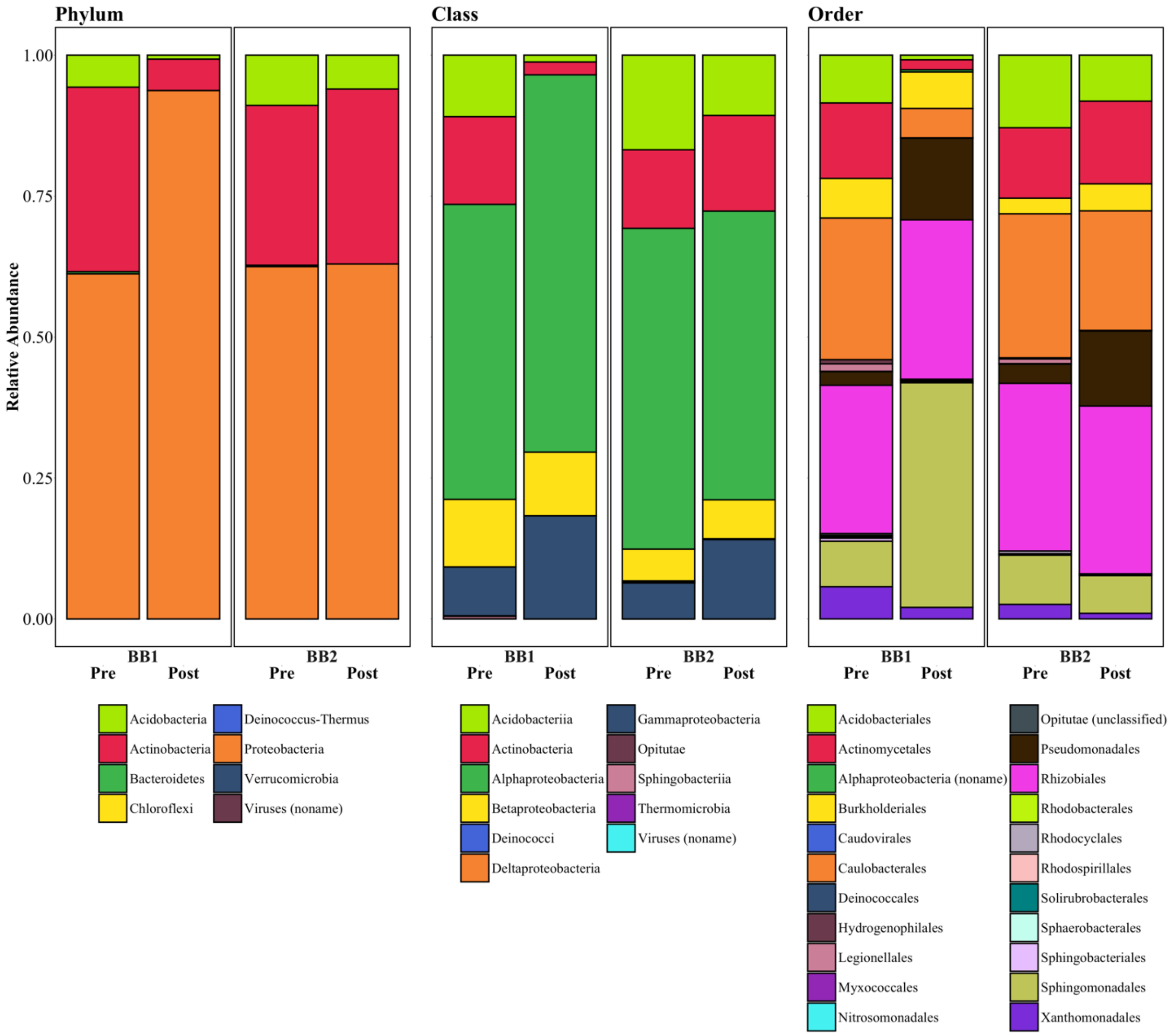


**Figure S5:** The relative abundance of the phylum, class, and order taxonomic levels. Values are averaged between the three sampling replicates; across all samples, there were 8 different phyla, 10 different classes, and 22 different orders identified. *BB (Biobed), *Pre - pre-pesticide application, *Post - post-pesticide application.

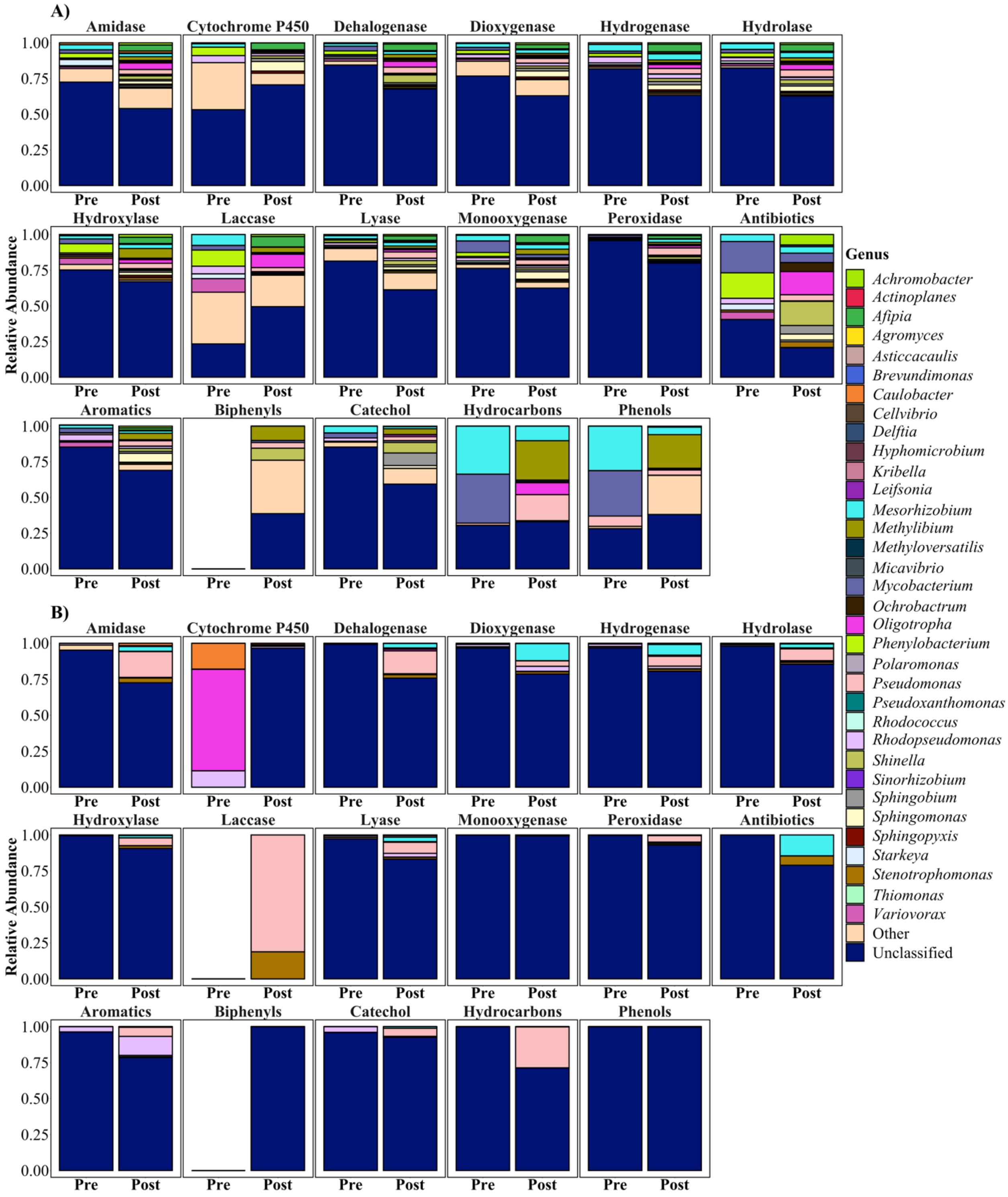


**Figure S6:** The relative abundance of the aromatic-degrading genes from Figure 3 in Biobed 1 from the A) metagenome and B) metatranscriptome, according to genus level. Genera displayed are those identified as have > 0.01 relative abundance (Figure 1A), significantly changing in abundance post-pesticide (Figure 1B), or being in substantial abundance (> 5.0 %) within the metatranscriptome. For the metagenome, abundances were averaged between the three replicates. *Pre - pre-pesticide application, *Post - post-pesticide application.
